# Enriching hippocampal memory function in older adults through video games

**DOI:** 10.1101/587006

**Authors:** Gregory D. Clemenson, Shauna M. Stark, Samantha M. Rutledge, Craig E.L. Stark

**Author notes:** To whom correspondence should be addressed at: 1424 Biological Sciences III, University of California, Irvine, Irvine CA, 92697-3800.

## Abstract

Healthy aging is accompanied by a steady cognitive decline with clear losses in memory. Animal studies have consistently demonstrated that simply modifying an animal’s living environment (known as environmental enrichment) can have a positive influence on age-related cognitive decline in the hippocampus. Previously, we showed that playing immersive 3D video games can improve hippocampal-based memory in young healthy adults, suggesting that the exploration of the large open worlds of modern-day video games may act as proxy for environmental enrichment in humans. Here, we replicated our previous video game study in older adults and showing that playing video games for 4 weeks can improve hippocampal-based memory in a population that is already experiencing age-related decline in this memory. Furthermore, we showed that the improvements last for up to 4 weeks past the intervention, highlighting the potential of video games as intervention for age-related cognitive decline.

## 1. Introduction

Healthy aging is closely associated with a reduction in hippocampal structure and function (Burke & Barnes, 2006; O’Shea, Cohen, Porges, Nissim, & Woods, 2016; S M Stark & Stark, 2017; Wilson, Gallagher, Eichenbaum, & Tanila, 2006). Despite this decline, studies in animals have shown that the aged hippocampus still retains a certain amount of plasticity that is susceptible to influence by the surrounding environment (Lee, Clemenson, & Gage, 2012). Environmental enrichment traditionally describes an experimental manipulation where an animal’s environment is enhanced to promote cognitive, physical, social, and other sensory stimulation (van Praag, Kempermann, & Gage, 2000). Despite the simplicity of this manipulation, environmental enrichment has been repeatedly shown to provide numerous structural and functional benefits to the hippocampus (Clemenson, Gage, & Stark, 2018). Importantly, environmental enrichment has been shown to improve both structural and cognitive deficits associated with aging, such as increased neurogenesis (Kempermann, Gast, & Gage, 2002; Leal-Galicia, Castañeda-Bueno, Quiroz-Baez, & Arias, 2008; Segovia, Yagüe, García-Verdugo, & Mora, 2006; Speisman et al., 2013), dendritic branching and spine density (Darmopil, Petanjek, Mohammed, & Bogdanović, 2009), expression of presynaptic proteins (Frick & Fernandez, 2003; Leal-Galicia et al., 2008; Saito et al., 1994), neurotransmitter release (Segovia et al., 2006), enhanced long-term potentiation and depression (Kumar, Rani, Tchigranova, Lee, & Foster, 2012; Stein, O’Dell, Funatsu, Zorumski, & Izumi, 2016), and related hippocampus-dependent behaviors (Kempermann et al., 2002; Segovia et al., 2006; Speisman et al., 2013). Whether the benefits of environmental enrichment are due to physical activity (Kobilo et al., 2011; Mustroph et al., 2012; van Praag, Kempermann, & Gage, 1999), spatial exploration (Freund et al., 2013), learning (Gould, Beylin, Tanapat, Reeves, & Shors, 1999; Leuner et al., 2004), or other aspects of the environment (Birch, McGarry, & Kelly, 2013; Clemenson et al., 2015; Steiner, Zurborg, Hörster, Fabel, & Kempermann, 2008), the surrounding environment can have a significant impact on the aging hippocampus of animals.

While the effects of environmental enrichment and its mechanisms are well defined in animals, it is less clear how this manipulation relates to humans and, specifically, to an aged population. Previously, we showed that playing immersive 3D video games could improve hippocampal-based memory in young adults (Clemenson & Stark, 2015). Given the long-standing relationship between the hippocampus and spatial memory (O’Keefe & Dostrovsky, 1971; Tolman, 1948) and the role of spatial exploration in environmental enrichment in animals (Freund et al., 2013), we hypothesized that the spatial exploration provided by the vast open worlds of modern day video games provide a human proxy for environmental enrichment. The goal of the present study was to determine if the improvements in hippocampal-based memory we observed with video game training in young adults would translate to an aged population, whom already experience age-related decline in hippocampal function.

We made several modifications to the current intervention to adapt to an older population. Similar to our previous study (Clemenson & Stark, 2015), participants played Angry Birds or Super Mario 3D World on a Nintendo Wii U. However, in place of a no-contact control, we employed an active control condition of playing Solitaire. Even though we did not observe an effect of playing Angry Birds in a younger population, we suspected that this condition might produce a benefit simply because the older adults in this study had no prior experience with modern video games or use a Nintendo Wii U. As we previously mentioned, one element of enrichment that has been described in the animal literature is the fact that enriched environments provide a certain amount of novelty and learning experiences to the animal that can influence the hippocampus (Gould et al., 1999; Leuner et al., 2004). Thus, there was a possibility that we would observe effects of learning within our Angry Birds group simply by virtue that playing any video game could provide a novel experience for them. For this reason, the Solitaire condition provided us with a control where participants actively played a game that they were familiar with. In addition, because this video game experience was so novel, we extended the amount of video game play from 2 to 4 weeks to address the steeper learning curve for this older population.

## 2. Materials and methods

### 2.1 Participants

We initially recruited 50 older adults to participate in our intervention. However, 5 participants were excluded prior to the analysis (3 scored more than 3 standard deviations below average on the recognition metric of the MST and 2 did not play the video game for the entire duration of the study). In total, 45 cognitively normal older adults (34 female, 11 male; mean age: 68.53 years, SD: 5.83; mean education: 16.31 years, SD: 2.29) were used for the analysis. Participants were recruited through local flyers and UCI affiliated groups, such as UCI MIND and the Alzheimer’s Disease Research Center (ADRC). While participants knew they would be playing video games, they were blind to our expectations based on the specific game. At the end of the study, all participants were compensated for their participation. All participants were screened for prior experience with modern video games using a modified version of the video game questionnaire used previously (Clemenson & Stark, 2015). Upon completion of a neuropsychological assessment, participants were randomly assigned to one of three intervention groups: Solitaire (6 female, 3 male; mean age: 68.7 years, SD 6.39; mean education: 17.11 years, SD: 1.05), Angry Birds (14 female, 4 male; mean age: 70.83 years, SD: 5.98; mean education: 15.94, SD: 2.65) or Super Mario (13 female, 5 male; mean age: 67.5 years, SD: 5.02; mean education: 16.2, SD: 2.3). All participants gave voluntary consent to participate and the study was conducted in compliance with the Institutional Review Board (IRB) of the University of California at Irvine.

### 2.2 Experimental Design

The entire intervention spanned eight weeks, including two neuropsychological assessments, four presentations of the Mnemonic Similarity Task (MST) using four distinct image sets, and four weeks of video game training (Figure 1A). Neuropsychological assessments occurred pre (week 0) and post video game training (end week 4). Administration of the MST occurred pre (week 0) and post video game training (end week 4), as well as during (end week 2) the video game training and four weeks after completion of the video game training (week 8), to assess any lasting effects of the intervention.

**Figure 1.**
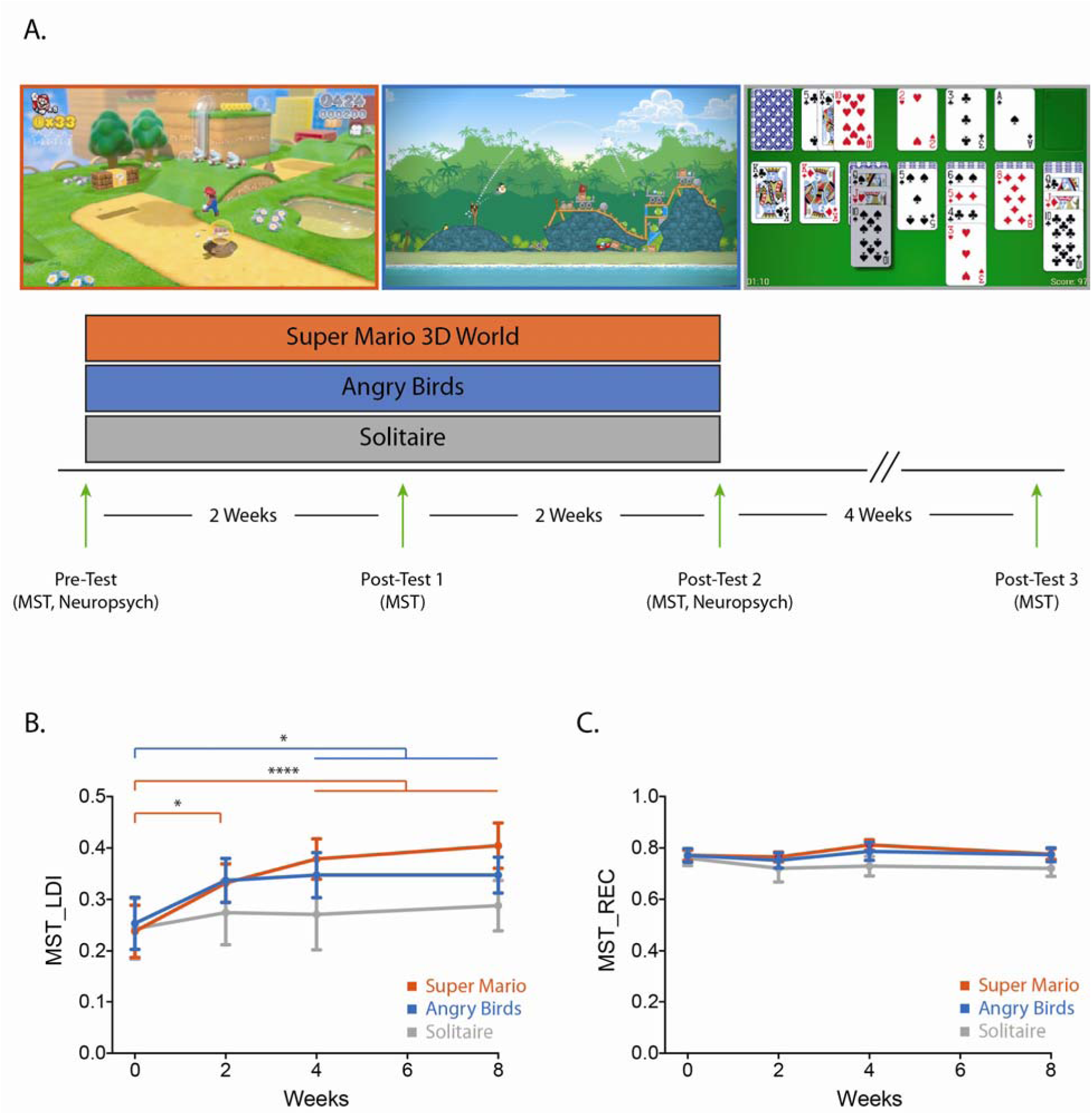
Experimental design and hippocampal-based memory performance. (A) Images of the three video games used and a schematic of the experimental design. (B) Performance in the hippocampal-mediated lure discrimination index (LDI) at pre-test (0-week) and post-test time points (2-week, 4-week, and 8-week), for all groups. (C) Performance in the general recognition memory (REC) at pre-test (0-week) and post-test (2-week, 4-week, and 8-week) time points, for all groups. All data are presented as mean ± SEM, *p < 0.05, ****p < 0.0001.

### 2.3 Neuropsychological assessment

All participants were given a neuropsychological assessment both prior-to and immediately following the video game intervention. The neuropsychological battery included tests for general cognition: Mini-Mental State Examination (MMSE) (Folstein, Folstein, & McHugh, 1975); memory: Rey Auditory Verbal Learning Test (RAVLT) (Rey, 1941) and Rey-Osterrieth (Rey-O) (Meyers & Meyers, 1995); executive functioning: Trails A & B (Tombaugh, 2004), Wechsler Adult Intelligence Scale (WAIS) Letter-Number sequence (Wechsler, 1997) and Stroop (Golden, 1978); and depression: Beck Depression Inventory (BDI) (A. Beck, 1972). We used two different word sets for the RAVLT and two different figure drawings for the Rey-O, pre and post intervention.

### 2.4 Mnemonic Similarity Task (MST)

The MST has previously been demonstrated to be sensitive to hippocampal function and age-related changes in the hippocampus (S. M. Stark, Yassa, Lacy, & Stark, 2013; Yassa, Muftuler, & Stark, 2010; Yassa, Stark, et al., 2010). The MST consists of two phases: a study and a test phase. In the first phase, participants were shown pictures of everyday objects and made simple indoor/outdoor judgement about each object (2s duration, ≥ 0.5s ISI). The second phase consists of a modified recognition test for the items previously seen or ones similar to those previously seen. Participants were shown either Targets (previously seen items), Lures (items similar to previously seen objects), or Novel foils (new items) and responded either ‘Old’, ‘Similar’, or ‘New’ respectively (2s duration, ≥ 0.5s ISI). In both phases, to better accommodate older participants, while the image disappeared from the screen after 2s, responses were self-paced. This allows for slower response times but prohibits excessive studying or scrutiny of the image that might lead to an altered strategy.

There are two metrics for assessing performance on the MST. The lure discrimination index (LDI) assessed mnemonic discrimination (or “behavioral pattern separation”) and is calculated as the probability of correctly identifying a ‘lure’ item as ‘similar’ minus incorrectly identifying a ‘foil’ item as ‘similar’ (lure|similar - new|similar). In addition to the LDI, a traditional recognition index (REC) serves as a measure of general recognition memory. REC is calculated as the probability of correctly identifying a ‘target’ item as ‘old’ minus incorrectly identifying a ‘new’ item as ‘old’ (target|old - foil|old). Importantly, the LDI metric is specifically sensitive to age-related changes in the hippocampus and cognitive decline whereas the REC metric is not (Kirwan et al., 2012; S. M. Stark, Stevenson, Wu, Rutledge, & Stark, 2015; S. M. Stark et al., 2013; Yassa & Stark, 2011).

### 2.5 Video game intervention

Participants were randomly assigned to train on one of three video games (Solitaire, Angry Birds, and Super Mario Bros 3D World) for 4 weeks, 30 minutes/day (max of 45 minutes/day). Participants were told to play at least 30 minutes/day (average time: 37.58 minutes/day, SD = 8.09 minutes). However, if at 30 minutes they were in the middle of a level/game, they were allowed to continue until completion of the level/game. Both Angry Birds and Super Mario 3D World groups played on the Nintendo Wii U, while Solitaire was played on the participants personal computer. Klondike Solitaire (GemMineMedia, https://www.gemmine.de/) was downloaded as an asset using the Unity 3D Engine (www.unity3d.com, Unity Technologies) and further modified (by G.D.C) to record the score and gameplay of the participants. All video game training occurred remotely at the participants’ homes. On the first day of video game training, an experimenter visited the residence of the participant to teach them how to set-up and play their respective game. In the case of Angry Birds and Super Mario, a Nintendo Wii U was provided to the participant and for Solitaire, a flash drive was brought and loaded onto a personal computer. If the participant did not have a TV for the Wii U, one was provided to them for the entire intervention (32-inch flat screen TV). Video game training time and performance was recorded by the participant, as well as by the Nintendo Wii U (Angry Birds and Super Mario) and the Solitaire game itself. All data was retrieved once the participant had completed the 4 weeks of video game training.

### 2.6 Statistical Analysis

All statistical analyses were performed using Prizm 7 (GraphPad, www.graphpad.com). Pre-planned comparisons were performed using 2-way ANOVAs with repeated measures and Sidak’s correction for multiple comparisons. A significance value of p<0.05 was used for all statistical analyses.

## 3. Results

### 3.1 Training on both Angry Birds or Super Mario 3D World led to improvements in hippocampal-based memory

Similar to our previous study (Clemenson & Stark, 2015), our primary question was whether playing Super Mario led to improvements in hippocampal-based memory. We entered the LDI metric of the MST into a repeated-measures 3×4 ANOVA to assess hippocampal-based memory across all three groups (AB, SM, Sol) and four time points (Pre, Post1, Post2, Post3). We found a significant main effect of time (Figure 1B; F(3, 126) = 9.33, p < 0.0001), with performance increasing over the course of the intervention, but no main effect of group (F(2,42) = 0.54, p = 0.58) or significant interaction (F(6,126) = 1.27, p = 0.27). A post-hoc analysis (Sidak correction for multiple comparisons at p < 0.05) revealed a significant improvement in the Super Mario group from the Pre-test to the 2-week (Post1), 4-week (Post 2), and 8-week (Post 3) time points, and a significant improvement in the Angry Birds group from the Pre-test to 4-week (Post 2) and 8-week 9Post 3) time points. There was no change from the Pre-test to any of the post-tests in the active-control Solitaire group. Importantly, general recognition memory (as measured by the REC metric of the MST) did not change across the intervention (Figure 1C; repeated measures 3×4 ANOVA main effect of time: F(3,126) = 1.463, p = 0.23), suggesting that the improvements we observed were specific to hippocampal-based memory. Unlike our previous study in young adults (Clemenson & Stark, 2015), where we found that improvements in the LDI metric correlated with how well participants performed in the video game, we did not find any such correlations between change in LDI from pre-test to post-test and video game performance (Solitaire: r^2^ = 0.01, p = 0.76; Angry Birds: r^2^ = 0.02, p = 0.64; Super Mario: r^2^ = 0.01 p = 0.74; data not shown).

To better understand the magnitude of these effects, we pooled all pre-test scores to compute the mean LDI of our entire pre-test sample (0.24) and then averaged all subsequent LDI post-test scores for each subject to compute our most robust estimate of the mean and standard deviation of the treatment effect for each experimental condition. The Solitaire group showed only a small, if any effect (LDI=0.28; Cohen’s d = 0.197), while the Angry Birds group showed a medium effect (LDI=0.34; d = 0.633) and the Super Mario group showed a large effect (LDI=0.37; d = 0.833).

### 3.2 Training on both Angry Birds or Super Mario 3D World led to an improvement in the Rey-Osterrieth complex figure task

While all three intervention groups performed similarly on the neuropsychological assessment across all measures at pre-test (Table 1), upon completion of the study, we found evidence that, like the LDI measure, performance on the Rey-O task improved in both the Angry Birds and Super Mario 3D world groups. Performance on the Rey-O involves making a copy of a complex figure while it is in front of the participant (Rey-O Copy) and drawing it again later from memory (Rey-O Delay). To account for any variation in drawing and copy performance and to isolate memory, we calculated the ratio between the Rey-O Delay score and the Rey-O Copy Score (Figure 2A). Comparing across all three intervention groups and two time points, we found a significant main effect of time (Figure 2A; 2×3 repeated-measures ANOVA: F(1,42) = 27.89, p < 0.0001), but no main effect of group (F(2,42) = 1.34, p = 0.27) or interaction (F(2,42) = 1.9, p = 0.16). However, a post-hoc analysis (Sidak correction for multiple comparisons at p < 0.05) revealed a highly significant improvement in both Angry Birds and Super Mario from pre-test to post-test (4-week) but not Solitaire. While both Angry Birds and Super Mario performed better than Solitaire at post-test (AB: p = 0.03 and SM: p = 0.04, uncorrected), this comparison did not survive corrections for multiple comparisons.

**Table 1.** This table presents the means and standard deviations of the neuropsychological tests used in this study at pre-test (0-week), across all groups.

| Pre-Test (mean, SD) | Solitaire | Angry Birds | Super Mario 3D World |
| --- | --- | --- | --- |
| MMSE | 29.44 (0.52) | 29.05 (0.93) | 28.77 (1.21) |
| RAVLT Total | 50.77 (10.07) | 51.72 (11.92) | 50.44 (12.01) |
| RAVLT Immediate | 10.22 (3.15) | 11.16 (3.14) | 10.5 (3.45) |
| RAVLT Delay | 9.66 (2.54) | 11.16 (3.16) | 11.27 (3.46) |
| Rey-O Copy | 35.11 (1.16) | 33.83 (2.53) | 33.61 (2.19) |
| Rey-O Delay | 15.5 (5.30) | 15.38 (6.46) | 16.63 (5.35) |
| Trails A | 35.33 (10.40) | 29.71 (12.24) | 27.49 (8.71) |
| Trails B | 60.25 (15.71) | 70.26 (21.80) | 71.30 (14.67) |
| Digit Span Total | 19.44 (4.06) | 17.94 (3.55) | 17.72 (4.22) |
| WAISIII LN Sequence | 18 (5) | 17.76 (5.6) | 16.94 (4.31) |
| Stroop Interference | 47 (7.56) | 48.94 (8.50) | 46.77 (9.73) |

**Figure 2.**
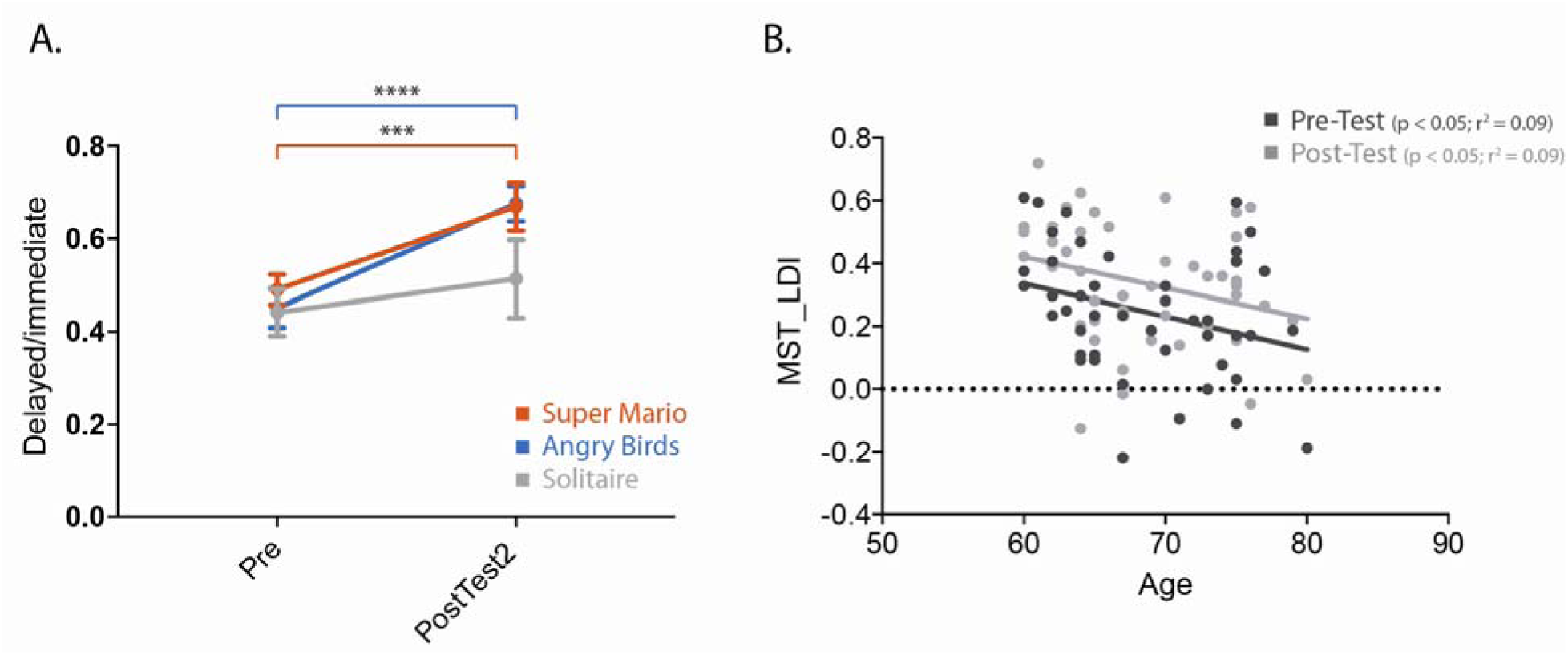
Performance on the Rey-O and change in hippocampal-based memory across age. (A) Performance on the Rey-O task at pre-test (0-week) and post-test (4-week) time points, for all groups. (B) Performance on the lure discrimination index (LDI) at pre-test (0-week) and post-test (4-week) time points for both Super Mario 3D World and Angry Birds groups combined, across age. All data are presented as mean ± SEM, ***p < 0.001, ****p < 0.0001.

### 3.3 The improvement in LDI was consistent across age

Previously, we showed that performance in the MST, and specifically the LDI metric, is sensitive to aging and continues to decline across age from 20-80 years of age (S. M. Stark et al., 2013). Here, in our more restricted range of older adults (60-80 years of age) that played Angry Birds and Super Mario 3D World, we again observed a significant decline in LDI with age (Figure 2B) at both pre-test (r^2^ = 0.09, p < 0.05) and post-test (r^2^ = 0.09, p < 0.05). These age effects were remarkably consistent in the pre-and post-test scores as the slopes were not reliably different from each other (F = 0.008, p = 0.92). In contrast, the pre-vs post-changes in LDI were reliably different (F = 5.305, p = 0.02). Furthermore, the change in LDI from pre-to post-test for both the Angry Birds and Super Mario groups did not correlate with age (r2 = 0.001, p = 0.84) suggesting that the benefits of the video game intervention on hippocampal-based memory are consistent across age.

## 4. Discussion

We tested the hypothesis that playing immersive 3D video games might act as a proxy for environmental enrichment and improve hippocampal-based memory in older adults. We found that older adults who participated in a novel video game intervention (both Super Mario 3D World and Angry Birds) for 4 weeks showed improved performance on an independent hippocampal-based memory task that persisted for up to four weeks after completion of the intervention. These findings, largely consistent with our previous study in young adults, highlight the potential value of video game play as an effective intervention for older adults.

In the current study, we found that, unlike our prior work in young adults (Clemenson & Stark, 2015), playing Angry Birds had a measurable impact on hippocampal-based memory. Although this improvement did not appear to be as robust as the Super Mario group and may have saturated early in the intervention, it did have a positive effect. One potential explanation is that the experience of console-based video game play itself is enriching for older adults. Our initial concern prior to this study was that because our older population had absolutely no experience with video games or video game consoles, showing up to their house with a large flat screen TV and a gaming console and having them play even a simple, novel video game may provide an enriching experience. As we mentioned previously, the novelty and learning experiences provided by enriched environments are thought to play an important role in its hippocampal influence (Gould et al., 1999; Leuner et al., 2004). Therefore, the benefits we observed here may be due to playing Angry Birds specifically, learning to use the Nintendo Wii U, or simply learning to use a joystick. Critically, actively playing a familiar game (on a familiar computer) did not have appear to have the same benefit.

Notably, the effects of video game training in our older adults persisted for up to four weeks after completion in the present study, whereas the improvement in our younger adults showed some evidence of regressing back towards baseline within two weeks of completion (Clemenson & Stark, 2015). There are several possible explanations for this difference. First, we should note that our prior work showed that after two weeks without gaming, the LDI scores were no longer reliably above pre-test baseline, but nor were they reliably below the immediately post training scores. That said, we would be remiss to not note that young college students are at a very different point in life than older adults. College is, arguably, a very enriching experience in adulthood. While video game play might be stimulating, there are countless other experiences happening in college (socially and educationally, both positive and negative) that may interfere or even replace the video game experience once it has stopped. Older adults, on the other hand, may have a more routine lifestyle and the video game intervention may represent a unique experience in itself. Another major difference between the two studies is the length of the intervention. Since the older adults played twice the amount of time as the younger adults (4 weeks instead of 2 weeks), it is possible that they reached or surpassed some threshold necessary for a longer-lasting change in hippocampal function. Further investigation is required to determine if this is due to the total number of hours played, consistent daily training, or some other aspect of the intervention.

In addition to improvements on the MST, we found that playing both Angry Birds and Super Mario, but not Solitaire, resulted in a significant improvement on the Rey-O. The Rey-O is a neuropsychological test used both clinically and in research to test several cognitive functions including attention, concentration, fine-motor coordination, visuospatial perception, nonverbal memory, and organizational skills (Shin, Park, Park, Seol, & Kwon, 2006). We examined Delay performance while controlling for the initial Rey-O Copy performance. The Rey-O Delay is highly sensitive to hippocampal amnesia (Rempel-Clower, Zola, Squire, & Amaral, 1996) and we have previously observed a positive correlation with LDI and hippocampal volume (Shauna M. Stark & Stark, 2017). Thus, it is not surprising that similar effects were observed for performance on the Rey-O and the MST’s LDI, both sensitive measures of hippocampal function. It is promising to see that the benefits from this video game intervention extend beyond the MST to other tasks and future studies should explore more sensitive measures of other cognitive domains. We should note that while the Rey-O has a long history and has its clear merits, repeat testing of it beyond two tests is difficult because alternative versions are not available. In contrast, the MST has at least 12 possible variants with no evidence of practice effects (Stark et al, 2015).

Lastly, the improvement on the MST (ΔLDI ≈ 0.1, d ≈ 0.8) observed here in our older adults is remarkably consistent with what we have observed previously in young adults using Super Mario 3D World (Clemenson & Stark, 2015), young adult competitive gamers specializing in different game genres (Clemenson & Stark, 2015), young adults using Minecraft to explore the world or learn to build complex structures (Clemenson, Henningfield, & Stark, 2019), and older adults in a real-world spatial memory intervention (Kolarik, Stark, & Stark, under review). In combination with our previous work in young adults and the fact that the observed improvement of older adults in this study did not change with age suggests that these interventions are equally effective across age ranges (18-22 and 60-80) and demonstrate the real potential of video games as an effective intervention resulting in improved memory function.

## 5. Conclusion

We showed that a four-week video game intervention in older adults can improve hippocampal-based memory and general cognitive behaviors. In conjunction with our previous work, we have shown that these video game interventions are practical, enjoyable, short (4 weeks) with a low daily commitment (~30 minutes/day), can be done remotely at home, and are highly effective. These studies highlight a real potential of using video games as a therapeutic intervention for age-related cognitive decline.

## Acknowledgements

This research was supported by a grant from the Dana Foundation. In addition, we would like to acknowledge support for participant recruitment from the UCI ADRC P50AG016573 and for additional salary support from NIA R21AG056145 and NIA R01AG034613. None of the authors have any conflicts of interest to declare.

